# Polymorphism analyses and protein modelling inform on functional specialization of Piwi clade genes in the arboviral vector *Aedes albopictus*

**DOI:** 10.1101/677062

**Authors:** Michele Marconcini, Luis Hernandez, Giuseppe Iovino, Vincent Houé, Federica Valerio, Umberto Palatini, Elisa Pischedda, Jacob Crawford, Rebeca Carballar-Lejarazu, Lino Ometto, Federico Forneris, Anna-Bella Failloux, Mariangela Bonizzoni

**Author notes:** University of California, Irvine, Department of Molecular Biology and Biochemistry, California, USA.

## Abstract

Current knowledge of the piRNA pathway is based mainly on studies on the model organism *Drosophila melanogaster*, where three proteins of the Piwi subclade of the Argonaute family interact with PIWI-interacting RNAs to silence transposable elements in gonadal tissues. In mosquito species that transmit epidemic arboviruses such as the Dengue and Chikungunya viruses, *Piwi* clade genes underwent expansion, are also expressed in the soma, and code for proteins that may elicit antiviral functions and crosstalk with other proteins of recognized antiviral mechanisms. These observations underline the importance of expanding our knowledge of the piRNA pathway beyond *D. melanogaster*.

Here we focus on the emerging arboviral vector *Aedes albopictus* and we couple traditional approaches of expression and adaptive evolution analyses with most current computational predictions of protein structure to study evolutionary divergence among Piwi clade proteins. Superposition of protein homology models indicate high structure similarity among all Piwi proteins, with high levels of amino acid conservation in the inner regions devoted to RNA binding. On the contrary, solvent-exposed surfaces showed low conservation, with several sites under positive selection. Expression profiles of *Piwi* transcripts during mosquito development and after infection with the Dengue 1 virus showed a concerted elicitation of all *Piwi* transcripts during viral dissemination, while the maintenance of infection primarily relied on the expression of *Piwi5*. In contrast, establishment of persistent infection by the Chikungunya virus is accompanied by an increased expression of all *Piwi* genes, particularly *Piwi4* and, again, *Piwi5*. Overall these results are consistent with functional specialization and a general antiviral role for *Piwi5*. Experimental evidences of sites under positive selection in *Piwi1/3*, *Piwi4* and *Piwi6*, further provide useful knowledge to design tailored functional experiments.

**Author summary:** Argonautes are ancient proteins involved in many cellular processes, including innate immunity. Early in eukaryote evolution, Argonautes separated into Ago and Piwi clades, which maintain a dynamic evolutionary history with frequent duplications and losses. The use of *Drosophila melanogaster* as a model organism proved fundamental to understand the function of Argonautes. However, recent studies showed that the patterns and observations made in *D. melanogaster*, including the number of Argonautes, their expression profile and their function, are a rarity among Dipterans.

In vectors of epidemic arboviruses such as Dengue and Chikungunya viruses, *Piwi* genes underwent expansion, are expressed in the soma, and some of them appear to have antiviral functions. Besides being an important basic question, the identification of which (and how) *Piwi* genes have antiviral functions may be used for the development of novel genetic-based strategies of vector control. Here we coupled population genetics models with computational predictions of protein structure and expression analyses to investigate the evolution and function of *Piwi* genes of the emerging vector *Aedes albopictus.* Our data support a general antiviral role for *Piwi5*. Instead, the detection of complex expression profiles with the presence of sites under positive selection in *Piwi1/3*, *Piwi4* and *Piwi6* requires tailored functional experiments to clarify their antiviral role.

## Introduction

First discovered for their role in plant developmental, proteins of the Argonaute family have been found in all domains of life, where they are essential for a wide variety of cellular processes, including innate immunity [1,2].

Recent studies provided evidence of evolutionary expansion and functional divergence of Argonautes in Dipterans, including examples in both the Ago and Piwi subclades [3]. Differences in function and copy number have also been found in other taxa such as nematodes [4], oomycetes [5] and higher plants [6], indicating a dynamic evolutionary history of this protein family. In eukaryotes, Argonautes are key components of the RNA interference (RNAi) mechanisms, which can be distinguished in small interfering RNA (siRNA), microRNA (miRNA) and the PIWI-interacting RNA (piRNA) pathways.

The siRNA pathway is the cornerstone of antiviral defense in insects. The canonical activity of this pathway is the Argonaute 2 (Ago2)-dependent cleavage of viral target sequences. Ago2 is guided to its target through an RNA-induced silencing complex (RISC) loaded with 21-nucleotide (nt)-long siRNAs. siRNAs are produced from viral double-strand RNAs intermediates by the RNAase-III endonuclease activity of Dicer-2 (Dcr2) and recognize the target based on sequence complementarity [7]. Dcr2 also possesses a DExD/H helicase domain that mediates the synthesis of viral DNA (vDNA) fragments [8], which appear to further modulate antiviral immunity [8]. vDNA fragments are synthetized in both circular and linear forms, in complex arrangements with sequences from retrotransposons, but details of their mode of action have not been elucidated yet [8,9]. We and others recently showed that the genomes of *Aedes spp.* mosquitoes harbor fragmented viral sequences, which are integrated next to transposon sequences, are enriched in piRNA clusters and produced PIWI-interacting RNAs (piRNAs) [10,11]. The similar organization between vDNAs and viral integrations, along with the production of piRNAs of viral origin (vpiRNAs) following arboviral infection of *Aedes* spp. mosquitoes, led to the hypothesis that the piRNA pathway functions cooperatively with the siRNA pathway in the acquisition of tolerance to infection [10,12,13].

Current knowledge on the piRNA pathway in insects is based mainly on studies on *Drosophila melanogaster*, where three proteins of the Piwi subclade, namely Argonaute-3 (AGO3), PIWI and Aubergine (AUB), interact with piRNAs to silence transposable elements (TEs) in gonadal tissues [14]. Interestingly, the piRNA pathway of *D. melanogaster* does not have antiviral activity and no viral integrations have been detected [15]. Additional differences exist between the piRNA pathway of *D. melanogaster* and that of mosquitoes, suggesting that *D. melanogaster* cannot be used as a model to unravel the molecular crosstalk between the siRNA and piRNA pathways leading to antiviral immunity in *Aedes* spp. mosquitoes. For instance, in *Aedes aegypti*, Piwi subclade has undergone expansion with seven proteins (i.e. Ago3, Piwi2, Piwi3, Piwi4, Piwi5, Piwi6 and Piwi7), which are alternatively expressed in somatic and germline cells and interact with both endogenous piRNAs and vpiRNAs [12,16,17]. Gonadal- or embryonic-specific expression is found for *Piwi1/3* and *Piwi7*, respectively [16], while *Ago3*, *Piwi4*, *Piwi5* and *Piwi6* are highly expressed in the soma and in the *Ae. aegypti* cell line Aag2 and contribute to the production of transposon-derived piRNAs [16,18]. Ago3 and Piwi5 also regulate biogenesis of piRNAs from the replication-dependent histone gene family [19]. Production of vpiRNAs is dependent on Piwi5 and Ago3 during infection of Aag2 cells with the *Alphavirus* CHIKV, Sindbis and Semliki Forest viruses (SFV), but relies also on Piwi6 following infection with the *Flavivirus* DENV2 [18,20–22]. Piwi4 does not bind piRNAs and its knock-down does not alter vpiRNA production upon infection of Aag2 cells with either SFV or DENV2 [18,23]. On the contrary Piwi4 coimmunoprecipitate with Ago2, Dcr2, Piwi5, Piwi6 and Ago3, suggesting a bridging role between the siRNA and piRNA pathways [21]. Despite these studies support an antiviral role for the piRNA pathway in *Aedes* spp. mosquitoes, a major challenge is to uncover the distinct physiological roles of Piwi proteins, if any. In duplicated genes, the presence of sites under positive selection is usually a sign of the acquisition of novel functions [24]. Additionally, under the “arm-race theory”, rapid evolution is expected for genes with immunity functions because their products should act against fast evolving viruses [25].

Besides being an important basic question, the understanding of functional divergence among Piwi proteins has applied perspectives for the development of novel genetic-based methods to control vector transmission. This is particularly relevant for mosquitos borne viruses, as several *Aedes* spp. species are expanding their spatial distribution and may contribute to disease outbreaks.

In recent years, the Asian tiger mosquito *Aedes albopictus* has emerged as a novel global arboviral threat, invading every continent except Antarctica from its native home range of South East Asia [26]. Because this species is a competent vector for a number of arboviruses such as chikungunya (CHIKV), dengue (DENV), yellow fever (YFV) and Zika (ZIKV) viruses, its establishment in temperate regions of the world fostered the re-emergence or the new introduction of these arboviruses [27]. For instance, Chikungunya outbreaks occurred in Italy in 2007 and 2017 [28,29]; France and Croatia suffered from autochthonous cases of Dengue and Chikungunya in several occasions since 2010 [30–33]; and dengue is re-emerging in some regions of the United States [34]. Despite its increasing public-health relevance, knowledge on *Ae. albopictus* biology and the molecular mechanisms underlying its competence to arboviruses are still limited in comparison to *Ae. aegypti*.

Here we elucidate the molecular organization, intraspecific polymorphism and expression of *Piwi* clade genes of *Ae. albopictus* in an evolutionary framework using a combination of molecular, population genomics and computational protein modelling approaches. We show that the genome of *Ae. albopictus* harbours seven *Piwi* genes, namely *Ago3*, *Piwi1/3*, *Piwi2*, *Piwi4*, *Piwi5*, *Piwi6* and *Piwi7*. For the first time in mosquitoes, we show sign of adaptive evolution in *Piwi1/3*, *Piwi4*, *Piwi5* and *Piwi6*, including sites in the MID and PAZ domains. Additionally, expression profiles during mosquito development and following infection with DENV or CHIKV support functional specialization of Piwi proteins, with a prominent and general antiviral role for the transcript of *Piwi5*.

## Results

### Seven *Piwi* genes are present in the genome of *Ae. albopictus*

Bioinformatic analyses of the current genome assemblies of *Ae. albopictus* (AaloF1) and the C6/36 cell line (canu_80X_arrow2.2), followed by copy number validation, confirmed the presence of seven *Piwi* genes (i.e. *Ago3*, *Piwi1/3*, *Piwi2*, *Piwi4*, *Piwi5*, *Piwi6* and *Piwi7*) in *Ae. albopictus* (S1 Table). Genomic DNA sequences were obtained for each exon-intron boundaries confirming in all *Piwi* genes the presence of the PAZ, MID and PIWI domains, the hallmarks of the Piwi subfamily of Argonaute proteins [35]. For *Ago3, Piwi1/3, Piwi2, Piwi4* and *Piwi6*, single transcript sequences that correspond to predictions based on the identified DNA sequences were retrieved (S1 Dataset). Sequencing results of the transcript from *Piwi5* showed a sequence 27 bp shorter than predicted on the reference genome, due to a 45bp gap followed by a 18b insertion, 110 and 333 bases after the ATG starting codon, respectively. This transcript still includes the PAZ, MID and PIWI domains. The presence of this transcript was further validated by northern-blot (Fig 1). For *Piwi7*, the transcript sequence also appears shorter than predicted (Fig 1). Alignment and phylogenetic analyses, in the context of currently annotated *Piwi* transcripts of Culicinae and Anophelinae mosquitoes, confirmed one-to-one orthologous pairing between *Ae. albopictus Piwi* gene transcripts and those of *Ae. aegypti* (S2 Table, S1 Fig). Interestingly, *Piwi5*, *Piwi6* and *Piwi7* transcripts group together and appear more similar to one of the two Aubergine-like transcripts annotated in different Anophelinae species than to *Aedes Piwi2*, *Piwi1/3* and *Piwi4* transcripts. Regarding the latter, *Piwi2* and *Piwi1/3* form a species-specific clade, rather than follow a speciation pattern. Independent duplication events in *Ae. aegypti* and *Ae. albopictus*, followed by convergent functional evolution, is unlikely if we consider the presence of orthologues in more distant species. Rather, the two genes, which based on *Ae. aegypti* chromosomal map on chromosome 1 and are ~20 kb apart [17], may be undergoing inter-locus gene conversion via nonreciprocal recombination, which result in between-loci homogenization.

**Figure 1.**
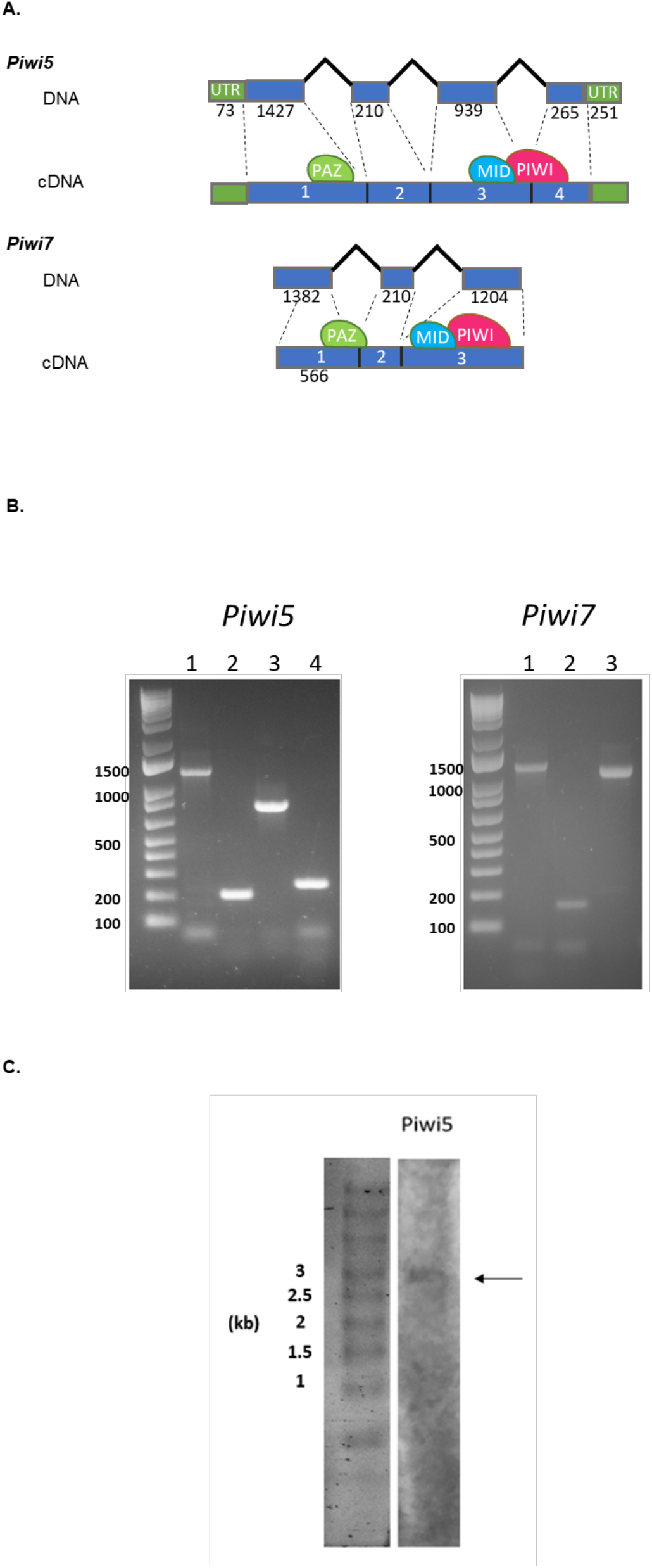
Gene and transcript structure of *Ae. albopictus Piwi5* and *Piwi7*. A) Schematic representation of the DNA structure of *Piwi5* and *Piwi7* genes and their corresponding transcripts as obtained from cDNA amplification of single sugar-fed mosquito samples. Exons and introns are shown by blue boxes and black lines, respectively, with corresponding length in nucleotide below each. The positions of the predicted PAZ, MID and PIWI domains are shown by green, blue and magenta ovals, respectively. Exon numbers correspond to lane numbers. B) Amplification of each exon of Piwi5 and Piwi7 on genomic DNA. Exon numbers correspond to lane numbers. C) Northern-blot results of *Piwi5* indicate the presence of a transcript of 3 kb.

### *Piwi* genes display high levels of polymorphism across populations and show signs of adaptive evolution

Across *Drosophila* phylogeny, genes of the piRNA pathway display elevated rates of adaptive evolution [36], with rapidly evolving residues not clustering at the RNA binding site, but being distributed across the proteins [3]. The RNA binding site is found within the PAZ domain, at the amino-terminal part of Piwi proteins [35,37]. The PIWI domain resides on the opposite side, at the carboxyl terminus. The PIWI domain belongs to the RNase H family of enzymes and the catalytic site is formed by three conserved amino acids (usually aspartate-aspartate-glutamate, DDE, or aspartate-aspartate-histidine, DDH) [35,38]. Between the PAZ and PIWI domains there is the MID domain. MID specifies strand- and nucleotide-biases of piRNAs, including their Uridine 5’ bias [39,40]. To evaluate the selective pressures acting along these genes, we analysed the polymorphism pattern in *Ae. albopictus* samples from wild-collected populations and from the Foshan reference strain. Synonymous and non-synonymous mutations were found for each gene in all populations (Fig 2), with *Piwi1/3* displaying the lowest polymorphism (Table 1).

**Table 1.** Polymorphism of *Aedes albopictus Piwi* genes in mosquitoes from the Foshan strain and wild-caught mosquitoes from La Reunion (Reu) and Mexico (Mex). We report the number of sequences (*n*), as well as the number of sites (*L*), segregating sites (*S*), polymorphism measured as π and θ, and the Tajima’s *D* statistic for both synonymous (s) and non-synonymous sites (a) for each gene and population (and for the pooled sample).

| | *n* | *L* | *L<sub>s</sub>* | *L<sub>a</sub>* | *S<sub>s</sub>* | *S<sub>a</sub>* | $\pi_s$ | $\pi_a$ | $\theta_s$ | $\theta_a$ | $\pi_a/\pi_s$ | *D<sub>s</sub>* | *D<sub>a</sub>* |
| --- | --- | --- | --- | --- | --- | --- | --- | --- | --- | --- | --- | --- | --- |
| ***Ago3*** |  |  |  |  |  |  |  |  |  |  |  |  |  |
| Pooled | 112 | 2832 | 680.2 | 2151.8 | 316 | 19 | 0.0699 | 0.0005 | 0.0878 | 0.0017 | 0.007 | -0.68 | -1.95 |
| Foshan | 32 | 2832 | 680.1 | 2151.9 | 124 | 5 | 0.0559 | 0.0004 | 0.0453 | 0.0006 | 0.007 | 0.89 | -0.82 |
| Mex | 48 | 2832 | 680.2 | 2151.8 | 253 | 14 | 0.0780 | 0.0007 | 0.0838 | 0.0015 | 0.009 | -0.25 | -1.60 |
| Reu | 32 | 2658 | 643.8 | 2014.2 | 189 | 4 | 0.0678 | 0.0002 | 0.0729 | 0.0005 | 0.003 | -0.27 | -1.50 |
| ***Piwi1-3*** |  |  |  |  |  |  |  |  |  |  |  |  |  |
| Pooled | 112 | 2658 | 644.3 | 2013.7 | 136 | 23 | 0.0319 | 0.0010 | 0.0399 | 0.0022 | 0.033 | -0.66 | -1.51 |
| Foshan | 32 | 2658 | 644.0 | 2014.0 | 10 | 2 | 0.0047 | 0.0003 | 0.0039 | 0.0002 | 0.064 | 0.68 | 0.44 |
| Mex | 48 | 2658 | 644.9 | 2013.1 | 117 | 21 | 0.0463 | 0.0017 | 0.0409 | 0.0024 | 0.037 | 0.48 | -0.89 |
| Reu | 32 | 2658 | 643.8 | 2014.2 | 52 | 4 | 0.0188 | 0.0004 | 0.0201 | 0.0005 | 0.021 | -0.23 | -0.48 |
| ***Piwi2*** |  |  |  |  |  |  |  |  |  |  |  |  |  |
| Pooled | 112 | 2625 | 644.0 | 1981.0 | 242 | 28 | 0.0760 | 0.0012 | 0.0710 | 0.0027 | 0.016 | 0.23 | -1.65 |
| Foshan | 32 | 2625 | 644.0 | 1981.0 | 115 | 10 | 0.0663 | 0.0017 | 0.0443 | 0.0013 | 0.026 | 1.88 | 1.11 |
| Mex | 48 | 2625 | 643.9 | 1981.1 | 184 | 15 | 0.0823 | 0.0010 | 0.0644 | 0.0017 | 0.012 | 1.01 | -1.28 |
| Reu | 32 | 2625 | 644.1 | 1980.9 | 151 | 6 | 0.0712 | 0.0005 | 0.0582 | 0.0008 | 0.007 | 0.85 | -0.94 |
| ***Piwi4*** |  |  |  |  |  |  |  |  |  |  |  |  |  |
| Pooled | 112 | 2592 | 620.0 | 1972.1 | 268 | 61 | 0.0729 | 0.0025 | 0.0817 | 0.0058 | 0.034 | -0.36 | -1.82 |
| Foshan | 32 | 2592 | 620.1 | 1971.9 | 122 | 18 | 0.0610 | 0.0009 | 0.0489 | 0.0023 | 0.015 | 0.94 | -2.05 |
| Mex | 48 | 2592 | 619.8 | 1972.2 | 181 | 41 | 0.0692 | 0.0035 | 0.0658 | 0.0047 | 0.051 | 0.19 | -0.87 |
| Reu | 32 | 2592 | 620.1 | 1971.9 | 161 | 45 | 0.0699 | 0.0029 | 0.0645 | 0.0057 | 0.041 | 0.32 | -1.79 |
| ***Piwi5*** |  |  |  |  |  |  |  |  |  |  |  |  |  |
| Pooled | 112 | 2745 | 653.1 | 2091.9 | 148 | 23 | 0.0457 | 0.0016 | 0.0428 | 0.0021 | 0.035 | 0.22 | -0.66 |
| Foshan | 32 | 2793 | 664.5 | 2128.5 | 58 | 8 | 0.0361 | 0.0018 | 0.0217 | 0.0009 | 0.050 | 2.47 | 2.78 |
| Mex | 48 | 2745 | 652.9 | 2092.1 | 137 | 13 | 0.0470 | 0.0017 | 0.0473 | 0.0014 | 0.036 | -0.02 | 0.65 |
| Reu | 32 | 2793 | 663.4 | 2129.6 | 89 | 6 | 0.0326 | 0.0008 | 0.0333 | 0.0007 | 0.025 | -0.08 | 0.40 |
| ***Piwi6*** |  |  |  |  |  |  |  |  |  |  |  |  |  |
| Pooled | 112 | 2661 | 649.0 | 2012.0 | 242 | 8 | 0.0805 | 0.0010 | 0.0705 | 0.0008 | 0.013 | 0.47 | 0.82 |
| Foshan | 32 | 2661 | 648.3 | 2012.8 | 92 | 3 | 0.0632 | 0.0001 | 0.0352 | 0.0004 | 0.002 | 2.99 | -1.69 |
| Mex | 48 | 2661 | 649.9 | 2011.1 | 213 | 7 | 0.0840 | 0.0001 | 0.0739 | 0.0008 | 0.001 | 0.50 | -2.33 |
| Reu | 32 | 2661 | 648.5 | 2012.5 | 163 | 4 | 0.0784 | 0.0001 | 0.0624 | 0.0005 | 0.001 | 0.98 | -2.01 |
| ***Piwi7*** |  |  |  |  |  |  |  |  |  |  |  |  |  |
| Pooled | 112 | 1977 | 469.8 | 1507.2 | 192 | 33 | 0.0877 | 0.0036 | 0.0772 | 0.0041 | 0.041 | 0.45 | -0.42 |
| Foshan | 32 | 1977 | 469.8 | 1507.2 | 118 | 15 | 0.0905 | 0.0034 | 0.0624 | 0.0025 | 0.038 | 1.71 | 1.25 |
| Mex | 48 | 1977 | 469.9 | 1507.1 | 150 | 23 | 0.0905 | 0.0034 | 0.0719 | 0.0034 | 0.038 | 0.93 | -0.04 |
| Reu | 32 | 1977 | 469.6 | 1507.5 | 137 | 17 | 0.0803 | 0.0030 | 0.0724 | 0.0028 | 0.037 | 0.41 | 0.24 |

**Figure 2.**
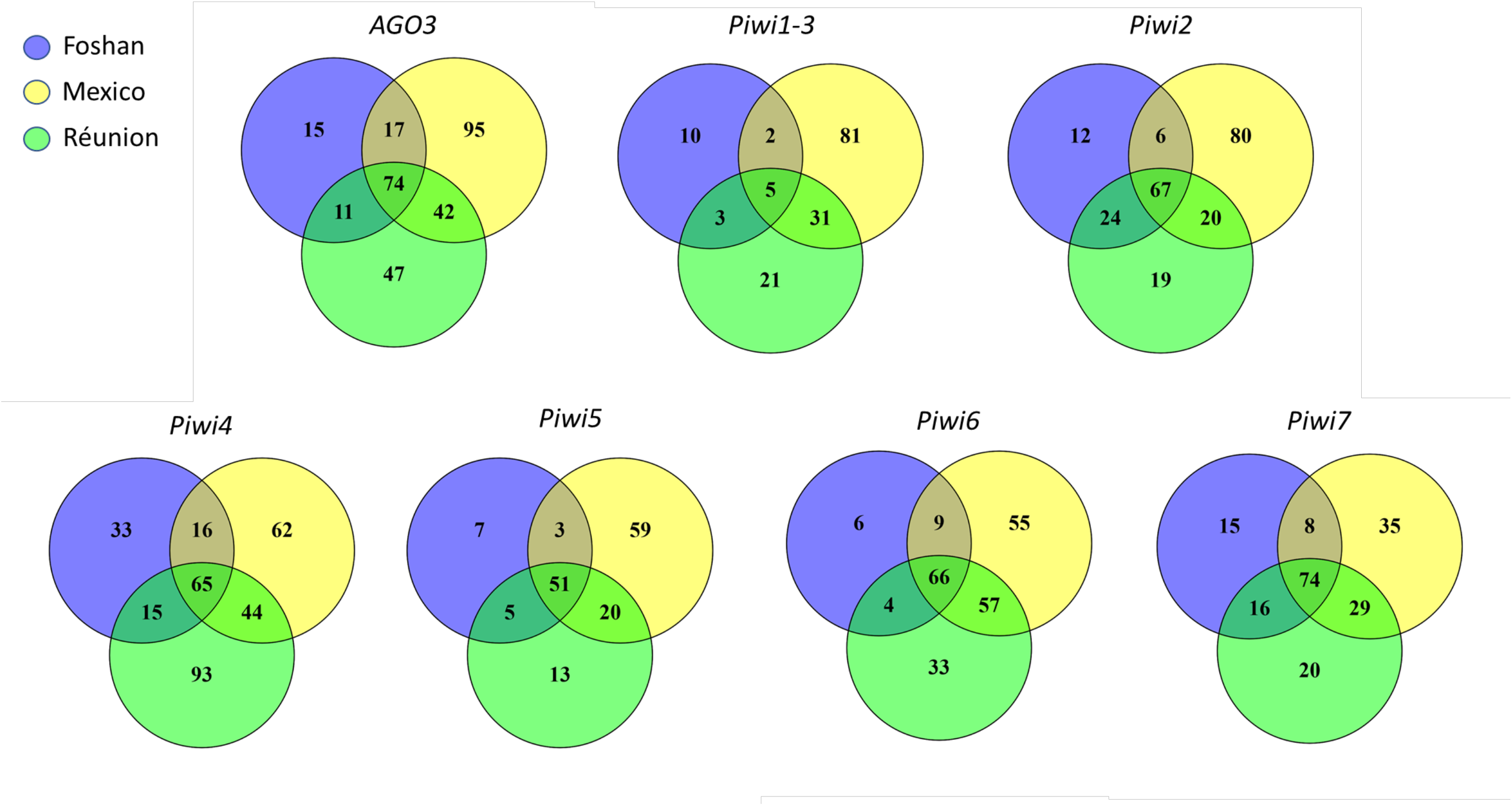
Venn diagrams showing the number of positions harbouring synonymous and non-synonymous mutations in tested samples for each *Piwi* gene.

As expected, the laboratory strain Foshan showed the lowest levels of variability and Tajima’s *D* values that contrast (in sign) from those of the other populations and from the pooled sample, consistent with a strong bottleneck associated to the strain establishment. In *Piwi4*, between 20 and 80 non-synonymous variants could be found inside and in proximity of the PAZ, MID and PIWI domains (S2 Fig.A), ten of these mutations were shared across all populations (S3 Table). The 5’ region of *Piwi5* harboured several indels: two in-frame variants (i.e. 94_99del; 113_118del) were shared across all populations and were present in homozygosity in at least one sample (S2 Fig.B), suggesting that they are not detrimental. *Ago3* and *Piwi6* have very low non-synonymous nucleotide diversity, suggesting strong constraints at the protein level. However, the McDonald-Kreitman test [41] found signatures of adaptive evolution in *Piwi1/3* and also in *Piwi6*, consistent with divergent positive selection followed by purifying selection (Table 2.A). In contrast, *Piwi4* has a significant deficit of non-synonymous substitutions and/or excess of polymorphic non-synonymous segregating sites (Table 2.A). In this gene, Tajima’s *D* is negative but in line with the values of the other *Piwi* genes, and the high non-synonymous polymorphism may reflect selection of intraspecific diversifying selection, as expected in genes involved in immunity. Because positive selection may have acted at the level of very few sites, this not contributing to the gene-level non-synonymous substitution pattern, we explicitly tested models of codon evolution. Signs of positive selection were found at different sites, including one site in the Linker2 and one site in the MID domain of *Piwi1/3*, two sites in the PAZ domain of *Piwi4*, two sites in the Flex domain of *Piwi5* and three sites, two in the Flex and one in the Linker2 domains, of *Piwi6* (Table 2.B). Haplotype reconstruction of our samples showed that these mutations can co-occur on the same gene, with the only exception of Y278D+H287P in Piwi4 and A67P+G86S in *Piwi6*.

**Table 2.** Insights into Evolutionary divergence of Piwi genes in *Ae. albopictus*. A) McDonald-Kreitman test for each *Piwi* gene using the orthologous sequences of *Ae. aegypti* as outgroup. NI = Neutrality Index; Alpha = proportion of base substitutions fixed by natural selection; *P* estimated using Fisher’s exact test. B) Output of Codeml with significant results

| A. McDonald-Kreitman test |  |  |  |  |  |  |  |
| --- | --- | --- | --- | --- | --- | --- | --- |
|  | *Ago3* | *Piwi1/3* | *Piwi2* | *Piwi4* | *Piwi5* | *Piwi6* | *Piwi7* |
| **NI** | 0.582 | 0.516 | 0.9 | 3.888 | 0.696 | 0.154 | 0.745 |
| **alpha** | 0.418 | 0.484 | 0.1 | -2.888 | 0.304 | 0.846 | 0.255 |
| **P** | 0.114 | 0.008 | 0.785 | < 0.001 | 0.18 | < 0.001 | 0.272 |

regarding sites under positive selection.
| B. Codeml output for sites under positive selection |  |  |  |  |
| --- | --- | --- | --- | --- |
| Gene | Position<sup>1</sup> | Reference>Mutant<sup>2</sup> | P<sup>3</sup> | Domain<sup>4</sup> |
| *AGO3* | - | - | - |  |
| *Piwi1-3* | 485 | K>R | 0.965* | Linker2 |
|  | 548 | M>I | 0.984* | MID |
| *Piwi2* | - | - | - |  |
| *Piwi4* | 278 | Y>D | 0.993** | PAZ |
|  | 287 | H>A,D,P,V | 1.000** | PAZ |
| *Piwi5* | 89-90 | SA>PT | 1.000* | Flex |
|  | 139 | T>A | 1.000* | Flex |
| *Piwi6* | 67 | A>P | 0.992** | Flex |
|  | 86 | G>R,S | 0.957* | Flex |
|  | 258 | V>I | 0.999** | Linker2 |
| *Piwi7* | - | - | - |  |

<sup>1</sup> sites where signs of positive selection ( $\omega > 1$ ) were found; <sup>2</sup> reference amino acid and alternative missense variant; <sup>3</sup>probability that $\omega > 1$ under the Bayes empirical Bayes (BEB) method (\* = $P > 0.95$ ; \*\* = $P > 0.99$ ); <sup>4</sup>protein domain based on computational
predictions of molecular structures. Domains are as follows: Linker2, linker region between PAZ and MID; PAZ domain; MID domain; and Flex, the Flexible stretch at the N-terminus.

Finally, to gain insight on how variable *Piwi* genes are in comparison to slow- and fast-evolving genes of *Ae. albopictus*, we collected variability data of sets of genes previously identified to have slow and high evolutionary rates [42]. For each population, we compared the overall level of polymorphism (LoP) of the *Piwi* genes and of a dataset of fast-evolving genes (FGs) to that measured for a dataset of slow-evolving genes (SGs) (Pischedda et al., 2019). Our results indicate that *Piwi4*, *Piwi6* and *Piwi7* have LoP values comparable to those of FGs, while *Ago3* and *Piwi5* do not significantly deviate from the LoP values of SGs. Piwi1/3 appears to be conserved (Fig 3).

**Figure 3.**
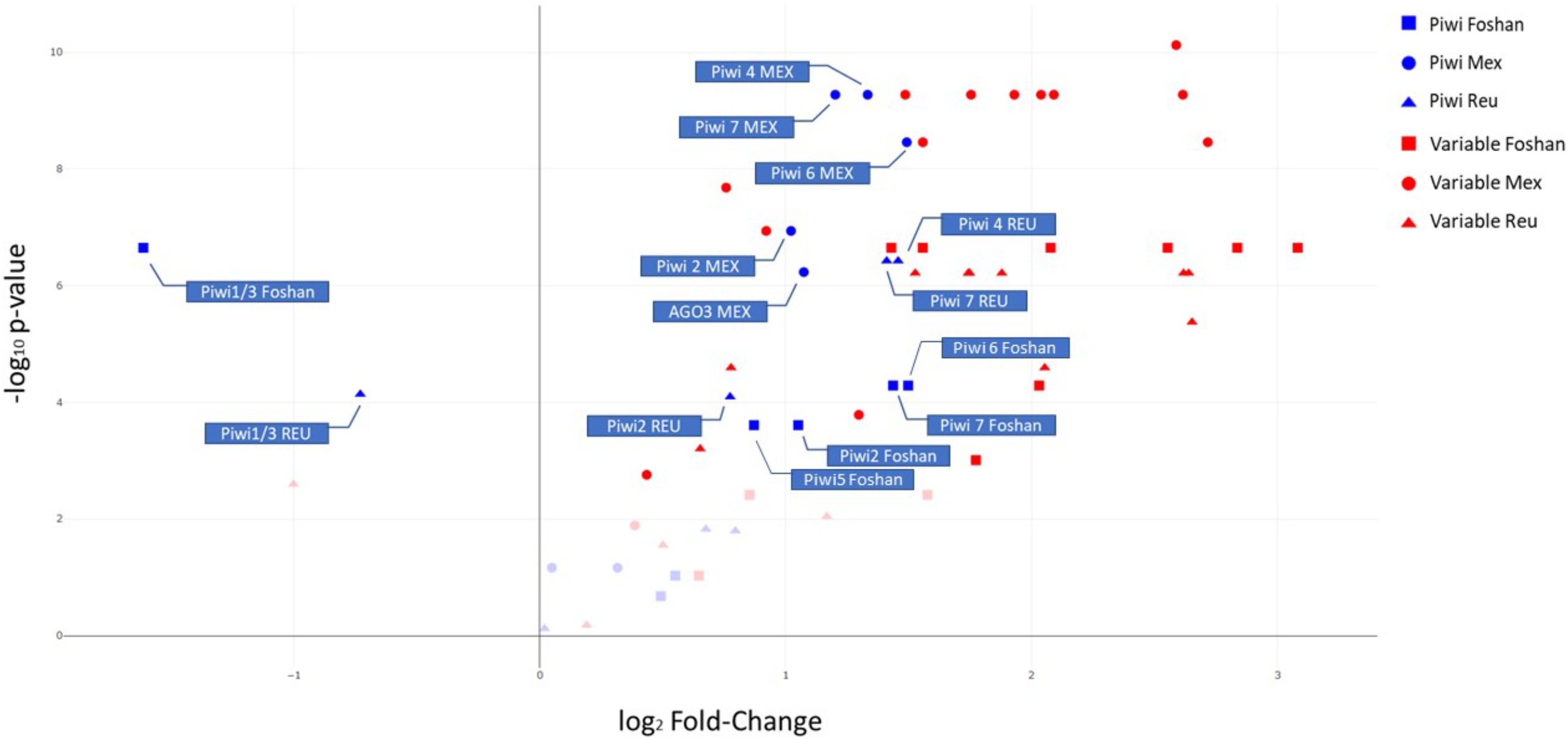
Volcano plot. Level of polymorphism (LoP) comparison between slow-evolving genes (SGs), fast-evolving genes (FGs) and *Piwi* genes by population. Genes on the right side of the panel have LoP values greater than those of SGs, while genes on the left side have LoPs smaller than SGs. The y-axis represents the −log10 p-values of the Kolmogorov-Smirnov test. Faint datapoints are not significant after Bonferroni correction for multiple testing (−log10 0.0024 (0.05/21 genes) = 2.62).

### Computational predictions of molecular structures

The functional significance of the mutations under selection, as well as that of all the shared missense mutations in the PAZ and PIWI domains, was tested by computing predictions of three-dimensional molecular structures of the Piwi proteins using the most-recent X-ray crystallography structure of Argonaute proteins as templates [43,44]. Homology modelling revealed high structural conservation among the seven Piwi proteins despite sequence heterogeneity (S2 Fig.; Fig 4.A).

**Figure 4.**
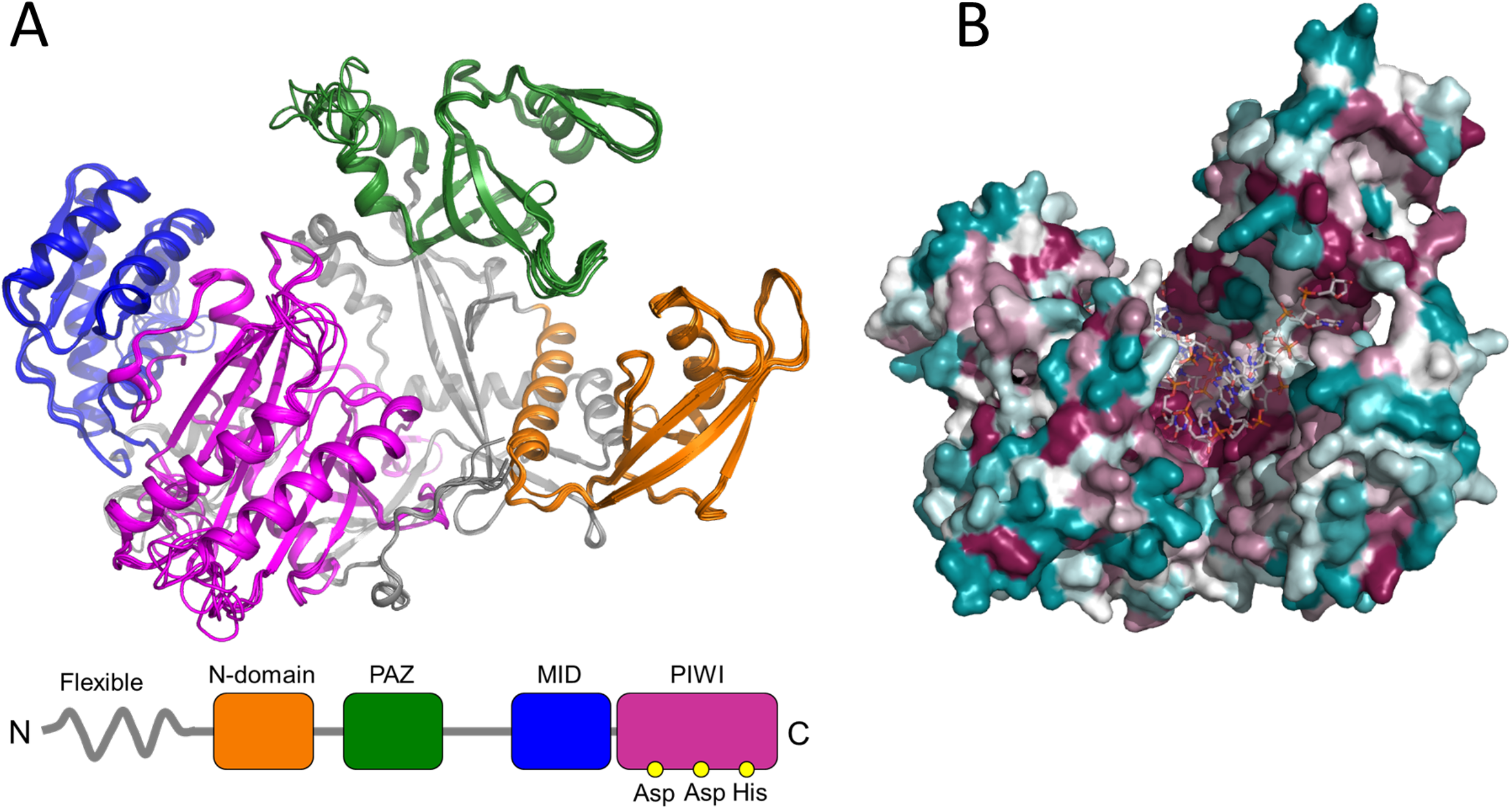
Computational homology models of the *Ae. Albopictus Piwi* proteins. Homology models were generated for the seven *Piwi* genes as described in the methods section. A) Superposition of cartoon representations of Piwi homology models, with highlight of domain organization: the N-terminal domain is shown in orange, the PAZ domain in green, the MID domain in blue and the PIWI domain in magenta. B) *CONSURF* [91]overview of the amino acid sequence conservation mapped on three-dimensional homology models in a putative RNA-bound arrangement based on the structure of human Argonaute bound to a target RNA (PDB ID 4Z4D), coloured from teal (very low conservation) to dark magenta (highly conserved).

Similarly, to *D. melanogaster*, the highest levels of amino acid sequence conservation were found in the regions that, based on homology modelling, define the inner pocket of Argonaute molecular assembly where the RNA binds. Significantly lower sequence conservation was found on the proteins surface (Fig 4.B). Based on our computational predictions, we could not detect aminoacidic polymorphisms that would affect RNA binding or processing, suggesting that all *Ae. albopictus* Piwi proteins may retain the Argonaute-like functions. Mapping of mutations under positive selection (Table 2.B) on the homology models showed that all variant amino acids were in regions distant from the predicted RNA-binding and/or processing sites, suggesting that these mutations are unlikely to alter protein folding, but could influence its stability.

### Developmental profile of *Ae. albopictus Piwi* genes

To further gain insights on the functional specialization of *Piwi* genes, we assessed their expression profile throughout mosquito development, namely at 4-8 hours (h) after deposition to capture the maternal-zygotic transition in expression, at late embryogenesis (i.e. 12-16 h and 16-24 h post deposition), at two time points during larval development (i.e. 1^st^ and 4^th^ instar larvae) and at pupal and adult stages (for the latter only we sampled separately males and females). Adult females were dissected to extract ovaries from the carcasses both from females kept on a sugar diet and 48 h after a blood meal, when a peak in *Piwi* gene expression was previously observed [45].

Expression levels of *Ago3*, *Piwi4*, *Piwi5*, *Piwi6* and *Piwi7* are at their peak in the embryonic stages, although at different time points (Fig 5). Overall, *AGO3*, *Piwi1/3, Piwi2* and *Piwi6* have a similar trend during development showing a second peak of expression in adult females and their ovaries, while the expression levels of *Piwi4*, *Piwi5* and *Piwi7* remain constant. In details, *Piwi7* is mostly expressed 4-8h after deposition*, Piwi5* and *Piwi6* are mostly expressed after 8-16h and *Ago3* and *Piwi4* have their pick of expression at 16-24h. On the contrary, *Piwi1/3* and *Piwi2* are mostly expressed in ovaries extracted from blood-fed and sugar-fed females, respectively (Fig 5.A, S4.A table). Interestingly, when considering the absolute expression levels, *Piwi7* is the least expressed of all the genes at any tested time point, with limited expression seen only in embryos within 24 hours post deposition (i.e. Ct values for *Piwi7* ranged from 24.04 to 30.65, at 4-8h and 1^st^ instar larvae, respectively; at the same time points, Ct values for *AGO3* were 27.45 and 26.96.). These results are consistent with lack of expression from published RNA-seq data from adult mosquitoes.

**Figure 5.**
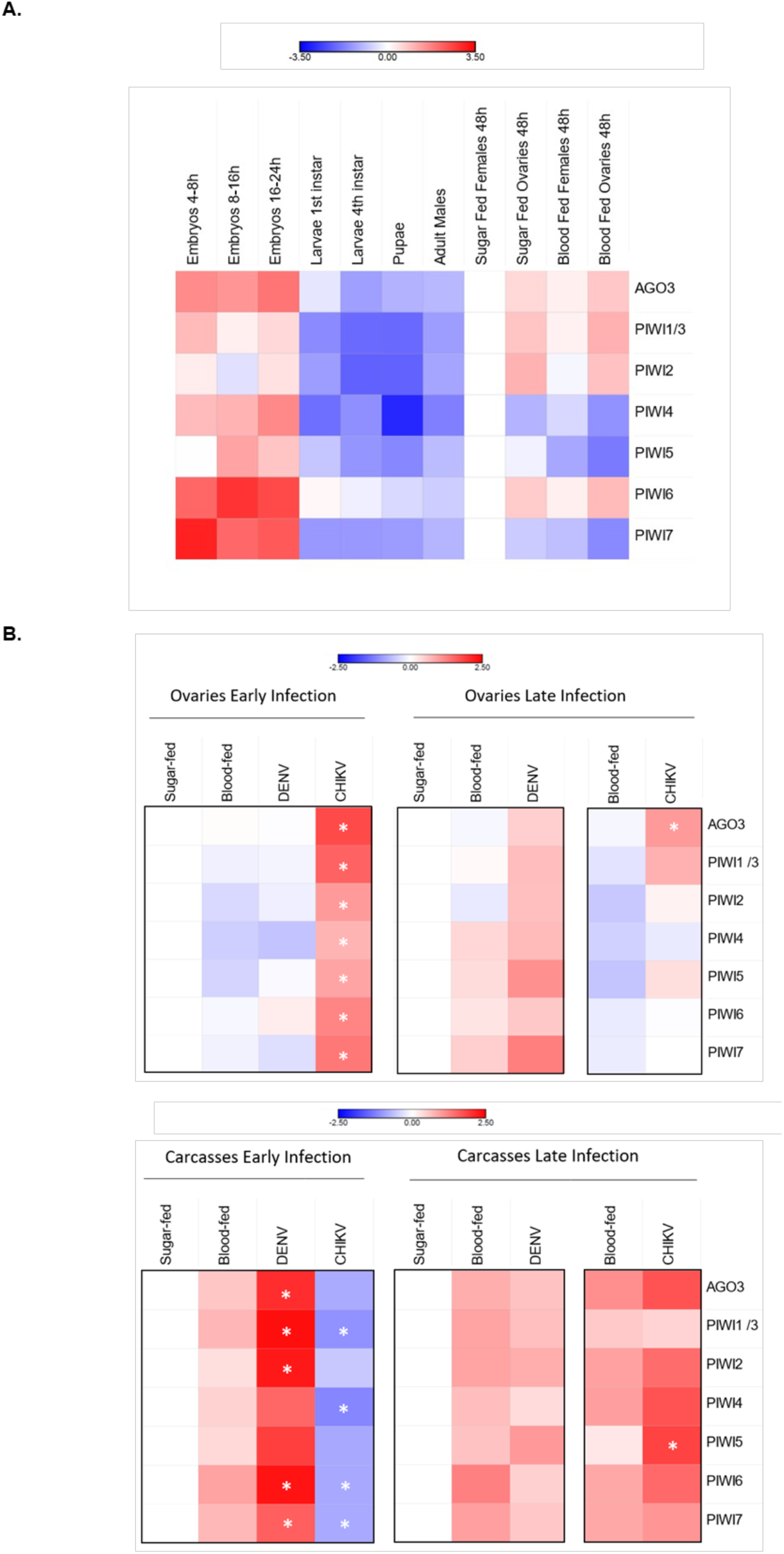
Expression profile of *Piwi* genes. Heatmap representations of log10 transformed fold-change expression values of each *Piwi* gene. A) Developmental expression pattern of the *Piwi* genes normalized on expression in early embryos (4-8h). B) Expression pattern of *Piwi* genes following viral infection normalized with respect to sugar-fed samples. Expression was verified in ovaries and carcasses separately, during the early and late stages of infections, that is 4 days post infection for both viruses and 14 or 21 dpi for CHIKV and DENV, respectively. Each day post infection was analysed with respect to sugar and blood-fed controls of the same day. * indicates significant difference (P<0.05) between infected samples and the corresponding blood-fed control.

Overall, at the adult stages, *Ago3* and all *Piwi* genes were more expressed in females than males. Expression in ovaries was higher than in the corresponding carcasses, in both sugar- and blood-fed females. Differences in carcasses vs. ovaries expression were more pronounced after blood-meal for *Ago3*, *Piwi1/3* and *Piwi6*, while expression of *Piwi2* was doubled in sugar-fed *vs.* blood-fed ovaries.

### *Piwi* genes expression following viral infection

Finally, we assessed whether the expression pattern of *Piwi* genes was altered upon DENV and CHIKV infection (Fig 5.B). Clear differences in the expression pattern of *Piwi* genes was seen both when comparing data from CHIKV-*versus* DENV-infected samples, and carcasses *versus* ovaries. In ovaries, during CHIKV infection all *Piwi* genes were significantly up-regulated compared to both sugar- and blood-fed mosquitoes. Four days post infection (dpi), the expression of *Ago3*, *Piwi1/3*, *Piwi6* and *Piwi7* was between 4 to 10 folds higher than that of *Piwi2*, *Piwi4* and *Piwi5*, which nevertheless were upregulated with respect to ovaries of sugar- and blood-fed mosquitoes. An opposite profile was seen in the carcasses, where all *Piwi* genes, particularly *Piwi1/3* and *Piwi4*, were down-regulated. At 4 dpi, CHIKV has already disseminated throughout the mosquito body, has reached the salivary glands and is able to be transmitted. CHIKV viral titer was reduced ten folds by 14 dpi and the profile of *Piwi* genes changed. Expression in the ovaries decreased between 3 (*Piwi5*) to 20 (*Piwi7*) times with respect to values observed at 4 dpi, but remained higher than the corresponding expression values in ovaries of both sugar and blood-fed mosquitoes. In carcasses all *Piwi* genes inverted their expression pattern during the infection phase, increasing up to more than 100 times in the case of *Piwi4*, *Piwi5* and *Piwi6.* At 14 dpi, expression of the *Piwi* genes was highest in CHIKV infected carcasses than in carcasses of sugar- and blood-fed mosquitoes.

For DENV, infection progresses differently than CHIKV. At 4 dpi there is no virus in the salivary gland, where the viral titer was measured at zero. By 21 dpi, DENV has established persistent infection [46]. At 4 dpi expression of *Piwi* genes was lower in DENV- and blood-fed ovaries than in ovaries of sugar-fed mosquitoes. The only exception was *Piwi6*, which was slightly up-regulated in ovaries of DENV-infected samples, but slightly down-regulated in ovaried of blood-fed mosquitoes. On the contrary, at the same time point, carcasses of DENV-infected samples showed a drastic increase in the expression of all *Piwi* genes with respect to blood-fed samples; this increase was between 7 to 87 times for *Piwi7* and *Piwi2*, respectively. By 21 dpi, expression in the ovaries increased for all *Piwi* genes, in comparison to what observed both at 4dpi and in blood-fed ovaries, suggesting the increase in expression of *Piwi* genes is related to DENV dissemination. Interestingly, if we compare levels of expression in CHIKV-infected ovaries at 4 dpi and DENV-infected samples at 21 dpi, corresponding to the time at which both viral species have disseminated throughout the mosquito body, we observe similar levels of fold-change expression of *Piwi4* and *Piwi7*, while *Ago3*, *Piwi1/3* and *Piwi6* show higher fold-change in CHIKV compared to DENV samples. Whether this trend is dependent on the viral species or viral titer requires further investigation. The same type of comparison in carcasses shows a higher fold-change expression level of all *Piwi* genes, particularly *Piwi1/3* and *Piwi5*, in DENV-*versus* CHIKV-infected samples, even if viral titer are lower for DENV (S4.B Table). Overall these results support the hypothesis of a concerted activity of all PIWI proteins during viral dissemination for DENV, and maintenance of infection rely on expression of primarily *Piwi5*. On the contrary, establishment of persistent CHIKV infection was accompanied by elicitation of all *Piwi* gene expression, particularly *Piwi4* and, again, *Piwi5*.

## Discussion

Recent experimental evidences extend the function of the piRNA pathway to antiviral immunity in *Aedes spp.* mosquitoes [12,15]. The broader roles of the piRNA pathway in *Aedes* spp. mosquitoes, compared to what is known in *D. melanogaster*, has been linked to the expansion and functional specialization of its core components [12,47,48]. Besides *Ago3*, the genome of *Ae. aegypti* harbours six *Piwi* genes (i.e. *Piwi1/3, Piwi2, Piwi4, Piwi5, Piwi6, Piwi7)*, some of which show tissue and development-specific expression profile and have been preferentially associated with either TE-derived or viral piRNAs [16,20,21]. These studies were based on the knowledge of the gene structure of each *Ae. aegypti Piwi* gene and the application of *ad hoc* RNAi-based silencing experiments and *in vitro* expression assays, but lacked an evolutionary perspective [18–21].

In this work we focused on the emerging arboviral vector *Ae. albopictus* and we show how the application of evolutionary and protein modelling techniques helps to unravel functional specialization of Piwi proteins. The genome of *Ae. albopictus* harbors one copy of *Ago3* and six *Piwi* genes (i.e. *Piwi1/3, Piwi2, Piwi4, Piwi5, Piwi6 and Piwi7*), each a one-to-one orthologue to the *Ae. aegypti Piwi* genes. The only exceptions are *Piwi2* and *Piwi1/3*, where the two genes from the same species cluster together. In *Ae. aegypti*, these two genes both map on Chromosome 1, separated by ~ 20kb, suggesting they may undergo frequent gene conversion.

All transcripts retain the PAZ and PIWI domains, which are the hallmarks of the Argonaute protein family [35]. By using homology modelling, we obtained predictions of molecular architectures for *Ae. albopictus Ago3* and *Piwi* proteins, onto which we mapped the putative boundaries of each domain. Superpositions and sequence comparisons allowed clear identification of the catalytic DDH triad within the PIWI domain of all modelled proteins. This conservation is consistent with strong sequence matching in the putative RNA binding regions of the PIWI, PAZ and MID domains and suggests the possible maintenance of slicer activity, albeit experimental validation of each isoform is necessary.

The expression of all *Piwi* genes was confirmed throughout the developmental stages and the adult life of the mosquito, both in ovaries and somatic tissues. Interestingly, *Piwi7* transcript expression starkly drops following early embryogenesis, to the point that we could detect it neither in RNA-seq analyses, nor in Northern-blot experiments (data not shown).

The expression of *Piwi* genes was elicited upon arboviral infection, indirectly confirming the antiviral role of the piRNA pathway. The expression profile of *Piwi* genes showed differences depending on both the species of infecting virus and on when the expression was measured. In CHIKV-infected samples, expression of *Piwi* genes was mostly elicited in ovaries or carcasses at 4 or 21 dpi, respectively. On the contrary, in DENV-infected samples, the highest expression of *Piwi* genes was seen in carcasses 4 dpi. These results are concordant with the timing in piRNAs accumulation following CHIKV or DENV infection. In *Ae. albopictus* mosquitoes infected with CHIKV, secondary piRNAs are not found 3 dpi, but are enriched 9 dpi [9]. In contrast, in *Ae. aegypti* mosquitoes infected with DENV2, piRNAs are the dominant small RNA populations 2 dpi [48]. Overall, these observations and our expression analyses support the hypothesis of an early activation of the piRNA pathway following DENV infection, but a late activation after CHIKV infection. Additionally, our expression analysis is consistent with a generalist antiviral role for Piwi5, which is elicited both during DENV and CHIKV infection [20], but suggest a more prominent role for Piwi6 and Piwi1/3 or Piwi4 and Ago3 during infection with DENV and CHIKV, respectively.

## Materials and methods

### Mosquitoes

*Aedes albopictus* mosquitoes of the Foshan strain were used in this study [10]. Mosquitoes were reared under constant conditions, at 28 °C and 70-80% relative humidity with a 12/12 h light/dark cycle. Larvae were reared in plastic containers at a controlled density to avoid competition for food. Food was provided daily in the form of fish food (Tetra Goldfish Gold Colour). Adults were kept in 30 cm^3^ cages and fed with cotton soaked in 0.2 g/ml sucrose as a carbohydrate source. Adult females were fed with defibrinated mutton blood (Biolife Italiana) using a Hemotek blood feeding apparatus. Mosquitoes from Mexico and La Reunion island were collected in 2017 as adults and maintained in ethanol 70% before shipment to Italy. All samples were processed at the University of Pavia.

### Mosquito infections

Foshan mosquitoes were infected with either DENV serotype 1, genotype 1806 or with CHIKV 06.21. DENV-1 (1806) was isolated from an autochthonous case from Nice, France in 2010 [49]. CHIKV 06-21 was isolated from a patient on La Reunion Island in 2005 [50]. Both strains were kindly provided by the French National Reference Center for Arboviruses at the Institut Pasteur. CHIKV 06-21 and DENV-1 1806 were passaged twice on cells to constitute the viral stocks for experimental infections of mosquitoes, on C6/36 cells for CHIKV 06-21 and on African green monkey kidney Vero cells for DENV-1 1806. Viral titers of stocks were estimated by serial dilutions and expressed in focus-forming units (FFU)/mL.

Four boxes containing 60 one-week-old females were exposed to an infectious blood-meal composed by 2 mL of washed rabbit red blood cells, 1 mL of viral suspension and 5 mM of ATP. The titer of the blood-meal was 10^7^ PFU/mL for CHIKV and 10^6.8^ PFU/mL for DENV. Fully engorged females were placed in cardboard boxes and fed with a 10% sucrose solution. Mosquitoes were incubated at 28 °C until analysis.

In parallel, mosquitoes were fed with uninfected blood-meal or kept on a sugar-diet and grown in the same conditions. Thirty mosquitoes were killed to be analyzed at days 4 and 14 post-infection (pi) for CHIKV, and at days 4 and 21 dpi for DENV.

To estimate transmission, saliva was collected from individual mosquitoes as described in [51]. After removing wings and legs from each mosquito, the proboscis was inserted into a 20 μL tip containing 5 μL of Fetal Bovine Serum (FBS) (Gibco, MA, USA). After 30 min, FBS containing saliva was expelled in 45 μL of Leibovitz L15 medium (Invitrogen, CA, USA) for titration. Transmission efficiency refers to the proportion of mosquitoes with infectious saliva among tested mosquitoes (which correspond to engorged mosquitoes at day 0 pi having survived until the day of examination). The number of infectious particles in saliva was estimated by focus fluorescent assay on C6/36 *Ae. albopictus* cells. Samples were serially diluted and inoculated into C6/36 cells in 96-well plates. After incubation at 28°C for 3 days (CHIKV) or 5 days (DENV), plates were stained using hyperimmune ascetic fluid specific to CHIKV or DENV-1 as primary antibody. A Fluorescein-conjugated goat anti-mouse was used as the second antibody (Biorad). Viral titers were 16,266±50,446 FFU and 155±125 FFU for CHIKV at 14 dpi and DENV at 21 dpi, respectively.

At the same time points mosquitoes that had been fed a not-infectious blood or kept on a sugar diet were sampled and dissected as above.

### Bioinformatic identification of *Piwi* genes in the *Ae. albopictus* genome

The sequences of the *Ae. aegypti Piwi* genes [52] were used as query to find orthologs in the reference genome of the *Ae. albopictus* Foshan strain (AaloF1 assembly) and in the genome of the *Ae. albopictus* C6/36 cell line (canu_80X_arrow2.2 assembly) using the BLAST tool in Vectorbase (www.vectorbase.org). Inferred coding sequences (CDS) where analysed in Prosite (Prosite.expasy.org/prosite.html) to screen for the typical PAZ and PIWI domains of Argonaute proteins [53].

### Copy number of *piwi* genes

qPCR reactions were performed using the QuantiNova SYBR Green PCR Kit (Qiagen) following the manufacturer’s instructions on an Eppendorf Mastercycler RealPlex4, on genomic DNA from four mosquitoes and using gene-specific primers, after having verified their efficiency (S5Table). DNA was extracted using DNA Isolation DNeasy Blood & Tissue Kit (Qiagen). Estimates of gene copy number were performed based on the *2*^−Δ*CT*^ method using *Piwi6* and the para sodium channel genes (AALF000723) as references [54].

### Structure of *Piwi* genes

DNA extracted from whole mosquitoes and dissected ovaries [55] was used as template in PCR amplifications to confirm the presence and the genome structure of each bioinformatically-identified *Piwi* gene. Primers were designed to amplify each exon, with particular attention to detect differences between paralogous *Piwi* genes (S1 Table). The DreamTaq Green PCR Master Mix (Thermo Scientific) was used for PCR reactions with the following parameter: 94 °C for 3 minutes, 40 cycles at 94 °C for 30 sec, 55 °C-62 °C for 40 sec, 72 °C for 1-2 minutes and final extension step of 72 °C for 10 minutes. PCR products were visualized under UV light after gel electrophoresis using 1-1.5% agarose gels stained with ethidium bromide and a 100 bp or 1 kb molecular marker. PCR products were either directly sequenced or cloned using the TOPO® TA Cloning® Kit strategy (Invitrogen) following the manufacturer’s instructions. DNA plasmids were purified using the QIAprep Spin Miniprep Kit and sequenced.

### *Piwi* gene transcript sequences and phylogeny

RNA was extracted using a standard TRIzol protocol from pools of 5 adult female mosquitoes to verify the transcript sequence of each *Piwi* gene. Sets of primers were designed for each gene to amplify its entire transcript sequence (S5 Table). PCR reactions were performed using a High Fidelity taq-polymerase (Platinum SuperFi DNA Polymerase, Invitrogen) following manufacturer’s instructions. PCR products were cloned using the TOPO® TA Cloning® Kit (Invitrogen) and plasmid DNA, purified using the QIAprep Spin Miniprep Kit, was sequenced. Rapid amplification of cDNA ends (RACE) PCRs were performed using FirstChoice RLM-RACE Kit (Thermo Fisher Scientific) to analyse 5’ and 3’ ends of the transcript sequences following manufacturer’s instructions. Amplification products were cloned and sequenced as previously indicated.

Sequences of the identified *Ae. albopictus Piwi* gene transcripts were aligned to sequences of *Culicidae* and *D. melanogaster Piwi* transcripts, as downloaded from VectorBase (www.vectorbase.org), using MUSCLE [56]. Maximum-likelihood based phylogenetic inference was based on RAxML after 1000 bootstrap resampling of the original dataset and was done through the CIPRESS portal (http://www.phylo.org/index.php/). Resulting tree was visualised using FigTree (http://tree.bio.ed.ac.uk/software/figtree/).

### Northern Blot analysis

10μg of total RNA from a pool of 10 sugar-fed females was run in a 1% × 2% agarose/formaldehyde gel (1 g agarose, 10 ml 10x MOPS buffer, 5.4 ml 37% formaldehyde, 84.6 ml DEPC water). Gels were washed twice in 20x SSC for 15 minutes prior to blotting. RNA was transferred to an Amersham Hybond-N+ nylon membrane (GE healthcare) using 20x SSC and cross-linked using UV light exposure for 1 minute. Probes were labelled with biotin using Biotin-High Prime (Roche). Hybridization and detection of biotinylated probes was performed using the North2South™ Chemiluminescent Hybridization and Detection Kit (Thermo Fisher Scientific) following manufacturer instructions.

### Polymorphisms of *Piwi* genes

We investigated *Piwi* gene polymorphism by looking at the distribution of single nucleotide polymorphism in whole genome sequence data from total of 56 mosquitoes, of which 24 from Mexico, 16 from the island of La Reunion island and 16 from the reference Foshan strain. Whole genome sequencing libraries were generated and sequenced on the Illumina HiSeqX platform at the Genomics Laboratory of Verily in South San Francisco, California to generate 150 basepair paired-end reads.

Illumina reads were mapped to *Piwi* gene transcript sequences using Burrows-Wheeler Aligner (BWA-MEM) [57] with custom parameters. Polymorphisms was tested by Freebayes [58]. Annotation of the detected mutations, as well counts of synonymous and non-synonymous variants, were performed in snpEff [59]. Frameshifts and non-synonymous variants were plotted using muts-needle-plot [60]. Venn diagrams of positions with mutations in the three tested samples were built using Venny 2.1 [61]. Haplotype reconstruction was performed using seqPHASE [62] and PHASE [63,64]. The inferred haplotypes were analysed with DnaSP [65], which estimated the number of segregating sites and the level of nucleotide diversity π [66] in both synonymous and non-synonymous sites. Based on the number of segregating sites and sample size, we also manually computed the nucleotide diversity estimator θ [67] and Tajima’s *D* statistic [68]. We also tested for signatures of adaptive evolution using the McDonald-Kreitman test [41], which compares the rate of polymorphism and substitutions in synonymous and non-synonymous sites. For this analysis we used alignments that included the orthologous sequences from *Ae. aegypti*.

Consensus sequences for each gene from each individual were also aligned in TranslatorX [69] using Clustalw [70] and used for Maximum-likelihood based phylogenetic inference based on RAxML after 1000 bootstrap under the GTRGAMMA model. Signs of selective pressure between populations [71] were investigated with Codeml in PAML v. 4.9 [72], as implemented in PAMLX [73]. In particular, we compared the M1a (nearly-neutral) model to the M2a (positive selection) model by inferring ω estimations and posterior probabilities under the Bayes empirical Bayes (BEB) approach [72].

The overall level of polymorphism (LoP) for slow-evolving genes (SGs) (AALF008224, AALF005886, AALF020750, AALF026109, AALF014156, AALF018476, AALF014287, AALF004102, AALF003606, AALF019476, AALF028431, AALF018378, AALF027761, AALF014448), fast-evolving genes (FGs) (AALF010748, AALF022019, AALF024551, AALF017064, AALF004733, AALF018679, AALF028390, AALF026991, AALF014993, AALF009493, AALF010877, AALF012271, AALF009839, AALF019413) and the *Piwi* genes was calculated for each population following the pipeline as in [42]. Briefly, SNPs and INDELs were inferred using four Variant callers (i.e. Freebayes [58], Platypus [74], Vardict [75] and GATK UnifiedGenotyper [76]) and the data merged and filtered with custom scripts. The LoP for each individual was calculated as the number of variants averaged over the region length, and the median value for each population was used for subsequent analyses. Statistical analyses were performed in R studio [77]. Fold-change differences were computed as the ratio of the median LoP for each Piwi gene and each FG gene over the median LoP of the SG genes. Statistical differences in LoP distribution was assessed via the Kolmogorov-Smirnov test and the p-value threshold was adjusted with the Bonferroni correction.

### Homology modelling

Computational structural investigations were carried out initially through the identification of the closest homologs based on sequence identity (using *NCBI Blast* [78]) and secondary structure matching (using *HHPRED* [79]). Homology model were then generated *MODELLER* [80] using on the structures *Kluyveromyces polysporus* Argonaute with a guide RNA (PDB ID 4F1N), Human Argonaute2 Bound to t1-G Target RNA (PDB ID 4z4d) [81], *T. thermophilus* Argonaute complexed with DNA guide strand and 19-nt RNA target strand (PDB ID 3HM9), and silkworm PIWI-clade Argonaute Siwi bound to piRNA (PDB ID 5GUH).

Computational models were manually adjusted through the removal of non-predictable N- and C-terminal flexible regions using *COOT* [82] followed by geometry idealization in *PHENIX* [83] to adjust the overall geometry. Final model quality was assessed by evaluating average bond lengths, bond angles, clashes, and Ramachandran statistics using Molprobity [84] and the *QMEAN* server [85] Structural figures were generated with *PyMol* [86].

### Developmental expression profile of *Piwi* genes

Publicly available RNA-seq data (runs: SRR458468, SRR458471, SRR1663685, SRR1663700, SRR1663754, SRR1663913, SRR1812887, SRR1812889, SRR1845684) were downloaded and aligned using Burrows-Wheeler Aligner (BWA-MEM) [57] to the current *Ae. albopictus* genome assembly (AaloF1). Aligned reads were visualized in Integrative Genomics Viewer (IGV) [87]. Total RNA was extracted from embryos, 1^st^ and 4^th^ instar larvae, pupae, and adults using Trizol (Thermo Fisher Scientific). Embryos consisted of two pools of 60 eggs at different time points after oviposition (i.e. 4-8 h, 8-16 h and 16-24 h). Adult samples consisted of males and females kept on a sugar-diet; females fed an uninfected blood-meal; and females fed a DENV-or CHIK-infected blood. All blood-fed females were dissected to separate ovaries from the carcasses. Females fed an uninfected blood-meal were sampled 48 h after blood-meal. These parameters were based on the results of previous studies on *Anopheles stephensi* and *Ae. aegypti* that showed high *Piwi* gene expression during early embryogenesis or 48-72 h post blood meal [45]. For each stage, RNA was extracted from pools of 10-15 mosquitoes, except for first instar larvae and embryos when 20 or 60 individuals were used, respectively.

RNA was DNaseI-treated (Sigma-Aldrich) and reverse-transcribed in a 20 μl reaction using the qScript cDNA SuperMix (Quantabio) following the manufacturer’s instructions. Quantitative RT-PCRs (qRT-PCR) were performed as previously described using two biological replicates per condition and the RPL34 gene as housekeeping [88]. Relative quantification of *Piwi* genes was determined using the software qBase+ (Biogazelle). Expression values were normalized with respect to those obtained from 4-8h embryos for the analysis of the developmental stages, and to sugar-fed females for the infection analyses.

### Expression analyses following infection

Fold-change expression values for each *Piwi* gene was assessed for non-infectious-blood-fed controls, CHIKV-infected and DENV-infected samples after normalization on sugar-fed controls. qRT-PCR experiments (Supplementary table 4) were set up for two replicate pools of 15 ovaries and 15 carcasses at days 4, 14 and 4, 21 for CHIKV and DENV, respectively and the corresponding sugar and non-infectious-blood controls. RNA extraction, qRT-PCR and data analyses were performed as described in the previous paragraph (see “Developmental expression profile of Piwi genes”). Fold-change differences significance was assessed using the Analysis of Variance (ANOVA) procedure [89,90] as implemented in qBASE+.

## Authors’ contributions

MM performed all experiments, analyzed the data and wrote the manuscript; LH performed PCR and qRT-PCR analyses and analyzed the data; GI contributed in identifying *Ae. albopictus Piwi* genes and their transcript sequences; VH contributed in infection experiments and analyzed the data; FV contributed in characterizing *Piwi* gene transcripts and their expression; UP, LO and EP contributed in bioinformatic analyses for *Piwi* gene polymorphism; AF supervised infection experiments analyzed data and wrote the manuscript; JC performed WGS of wild-caught mosquitoes and revised the manuscript; FF performed computational homology modelling and structural analyses, and revised the manuscript; RCL contributed in collecting wild mosquitoes and analyzed the data; MB conceived the study, analyzed the data and wrote the manuscript. All authors read and approved the final manuscript.

## Acknowledgments

We thank Monica Ruth Waghacore for insectary work.

## Supporting information

**S1 Table**. List of the core components of the piRNA pathway in *Ae. aegypti* and their orthologous in *Ae. albopictus*.

**S2 Table.** List of Transcript IDs and abbreviations of the Culicidae and Drosophilidae species included in the phylogenetic analyses.

**S3 Table.** Number of non-synonymous mutations found in mosquitoes of the Foshan strain (Foshan) and wild-caught samples from Mexico (Mex) and the island of La Reunion (Reu) divided by type (i.e. missense [M], frameshift [F], indel [I] and nonsense [N]) and number of sites in which

**S4 Table.** Relative expression values (log10 fold-change) of Piwi genes during development (A) and following viral infection (B) normalized with respect to sugar-fed samples. Samples (2 pools per condition, 15 individuals each) were analysed at 4 days post infection (early infection) and at 14 and 21 days post infection for CHIKV and DENV, respectively (late infection). Each condition was normalized to the corresponding Sugar-fed control and compared to the corresponding Blood-fed control. Ovaries and carcasses were analysed independently. * indicates statistically significant difference between infected and non-infected blood-fed samples (ANOVA framework). Relative expression values may mask differences in levels of expression. For instance, the Ct values of Piwi6, Piwi7 and Piwi1/3 in ovaries 4 days post infection with CHIKV were 30, 33.39 and 25.20, respectively. Ovaries of blood-fed samples at the same time point showed Ct values of 30.30, 33.93 and 26.55 for Piwi6, Piwi7 and Piwi1/3. When relative expression was calculated with respect to Ct values of RPL34, fold-changes in gene expression were comparable among the three genes in both conditions, but Ct values clearly indicate that Piwi7 is less expressed than both Piwi1/3 and Piwi6. These considerations were taken into account when describing results.

**S5 Table**. List of primers used for CDS analyses, copy number estimation, qPCR experiments and Northern Blot probe design.

**S1 Dataset.** CDS of the seven *Piwi* genes of *Ae. albopcitus*. The sequence of the **PAZ**, <u>MID</u> and ***PIWI*** domains is in **bold**, <u>underline</u> and ***bold-italics***, respectively.

**S1 Fig. Maximum likelihood cladogram generated from the alignment of transcript sequences of annotated *Piwi* genes in Culicinae.** Transcript IDs and species abbreviations are as listed in S2 Table. AlbPiwi3 is the same as Piwi1/3 in the text. *Piwi* gene transcripts from *Ae. albopictus* are in red, from *Ae. aegypti* in purple, from *Culex quinquefasciatus* in pink. Transcripts from *D. melanogaster Ago3*, *Piwi* and *Aubergine* genes are included for reference and shown in blue. All nodes were supported by bootstrap values higher than 50% with the exception of the three nodes with a black dot.

**S2 Fig. Polymorphism of *Piwi4* and *Piwi5*.** Lollipop plots representing position, amount and type of mutation along the coding sequences of *Piwi4* and *Piwi5* in mosquitoes of the Foshan strain, from la Reunion Island (Reu) and Mexico (Mex) as inferred by Freebayes and SnpEFF analyses. Only missense (blue), nonsense (red) and indels (orange) and frameshift (yellow) are shown. The PAZ, MID and PIWI domains are shown in green, blue and magenta, respectively. DDH residues positions are highlighted in the PIWI domain.

**S3 Fig.** Sequence alignment of *Aedes albopictus* Piwi proteins. Domain boundaries inferred from structural predictions are highlighted by coloured lines using the same colour coding as in figure 4 (Orange: N-terminus; Green: PAZ; Blue: MID; Magenta: PIWI). Conserved DDH residues found in PIWI are indicated by (▲). The “acc” line indicates relative solvent accessibility, ranging from blue (accessible) to white (buried). The sequence alignment was generated using EBI muscle [92] and depicted using ESPRIPT3 [93]

